# The pGinger family of expression plasmids

**DOI:** 10.1101/2023.01.23.524619

**Authors:** Allison N. Pearson, Mitchell G. Thompson, Liam D. Kirkpatrick, Cindy Ho, Khanh M. Vuu, Lucas M. Waldburger, Jay D. Keasling, Patrick M. Shih

**Affiliations:** Joint BioEnergy Institute, 5885 Hollis Street, Emeryville, CA 94608, USA; Biological Systems & Engineering Division, Lawrence Berkeley National Laboratory, Berkeley, CA 94720, USA; Department of Plant and Microbial Biology, University of California, Berkeley, CA 94720, USA; Environmental Genomics and Systems Biology Division, Lawrence Berkeley National Laboratory, Berkeley, California, USA; Department of Bioengineering, University of California, Berkeley, California, USA; Department of Chemical and Biomolecular Engineering, University of California, Berkeley, CA 94720, USA; The Novo Nordisk Foundation Center for Biosustainability, Technical University of Denmark, Denmark; Center for Synthetic Biochemistry, Institute for Synthetic Biology, Shenzhen Institutes for Advanced Technologies, Shenzhen, China; Innovative Genomics Institute, University of California, Berkeley, CA

## Abstract

The pGinger suite of expression plasmids comprises 43 plasmids that will enable precise constitutive and inducible gene expression in a wide range of gram-negative bacterial species. Constitutive vectors are composed of 16 synthetic constitutive promoters upstream of RFP, with a broad host range BBR1 origin and a kanamycin resistance marker. The family also has seven inducible systems (Jungle Express, Psal/NahR, Pm/XylS, Prha/RhaS, LacO1/LacI, LacUV5/LacI, and Ptet/TetR) controlling RFP expression on BBR1/kanamycin plasmid backbones. For four of these inducible systems (Jungle Express, Psal/NahR, LacO1/LacI, and Ptet/TetR), we created variants that utilize the RK2 origin and spectinomycin or gentamicin selection. Relevant RFP expression and growth data have been collected in the model bacterium *Escherichia coli* as well as *Pseudomonas putida*. All pGinger vectors are available via the Joint BioEnergy Institute (JBEI) Public Registry.

## Introduction

Precise and reliable control over gene expression is one of the most fundamental requirements of synthetic and molecular biology (1). Consequently, there has been considerable effort towards identifying myriad genetic elements that enable researchers to regulate the strength and timing of transcription across all domains of life (2–4). The end result of these efforts are often consolidated families of plasmid vectors that facilitate advanced genetic engineering, such as the BglBrick family of plasmids for *E. coli (5, 6)* and the jStack vectors used in multiple plant species (7). However, as the field of synthetic biology moves beyond traditional model organisms, families of expression vectors must be tailored to meet the specific requirements of particular hosts. Advances in non-model organisms often come in the form of species or genus specific toolkits (8–10), though more recently comprehensive plasmid toolkits have been developed and validated for a wide range of gram-negative organisms (11). Resources such as the Standard European Vector Architecture (SEVA) platform provide repositories of standardized sequences and constructs (12, 13). Still, given that many bacteria require very particular combinations of promoters, origins, and selectable markers to enable controlled gene expression, there remains a need for vectors that will allow rapid prototyping of genetic circuits in understudied bacteria.

To facilitate the exploration of non-model hosts, we have developed a small suite of plasmids that permit both constitutive and inducible expression from the broad host-range origin of replication BBR1 using a kanamycin selection marker. For a subset of the inducible systems that are known to work across multiple hosts, we have assembled combinatorial variants that utilize the compatible broad host-range origin RK2 (14) as well as both spectinomycin and gentamicin selection markers. This family of plasmids, which we have named the pGinger suite, requires no assembly of these parts, can be easily cloned into via standard Gibson assembly techniques, and has both digital sequences and physical samples that can be publicly accessed through the Joint BioEnergy Institute (JBEI) registry (15).

## Results

### Design and Architecture of pGinger Plasmids

All pGinger vectors express RFP with a consensus ribosomal binding site (RBS - TTTAAGAAGGAGATATACAT) derived from the BglBrick plasmid library. The overall conserved plasmid architecture and naming convention of the pGinger suite are shown in **Figure 1**.

**Figure 1:**
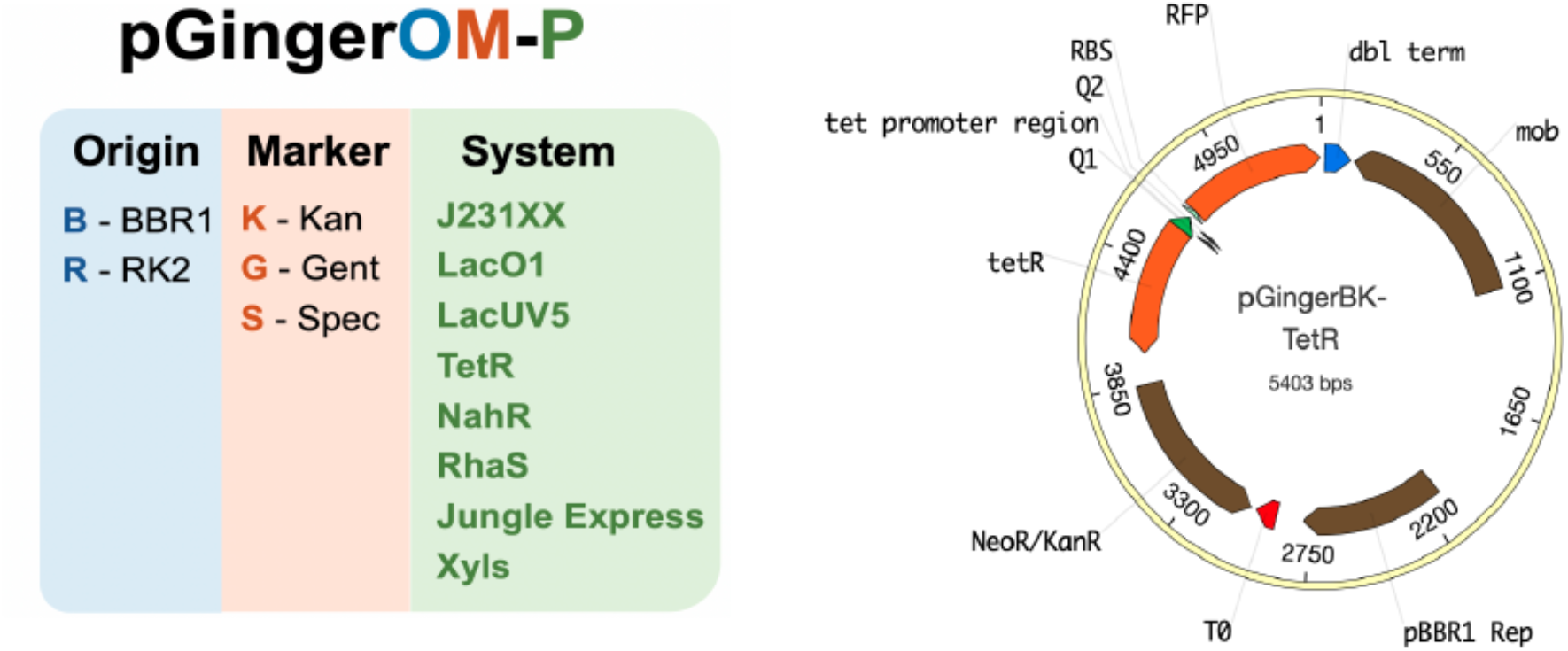
Plasmids architecture of the pGinger suite: The pGinger plasmids share a common naming convention where the first two letters after pGinger correspond to the origin and resistance marker respectively, followed by the expression system. All plasmids share the same architecture as the above map of pGingerBK-TetR, whereby a conserved RBS-RFP is downstream of the promoter followed by a strong terminator. All selectable markers are upstream of the promoter, with the origin between the marker and the RFP cassette.

The BBR1 origin and kanamycin cassette of relevant pGinger vectors were both derived from plasmid pBADTrfp (16). To develop a family of constitutive expression plasmids, the AraC coding sequence and promoter of pBADTrfp were replaced with 16 different synthetic promoters from the Anderson Promoter Library (http://parts.igem.org/Promoters/Catalog/Anderson). For the inducible vectors, the AraC coding sequence and promoter of pBADTrfp were replaced with the following seven inducible systems: Jungle Express - derived from pTR_sJExD-rfp (17); Psal/NahR - derived from pPS43 (18); Prha/RhaS - derived from pCV203 (18); Ptet/TetR - derived from pBbE2a-RFP (6); Pm/XylS - derived from pPS66 (18); LacO1/LacI - derived from pBbE6a-RFP (6); LacUV5/LacI - derived from pBbE5a-RFP (6). Three of these inducible systems, Pm/XylS, Psal/NahR, Prha/RhaS, utilize an activator or bifunctional transcription factor; the other systems feature transcriptional repressors. These BBR1 vectors contain the *mob* element that facilitates conjugal transfer. For four of the inducible systems (Jungle Express, Psal/NahR, LacO1/LacI, and Ptet/TetR), additional vectors were constructed that varied both the origin and antibiotic marker. All RK2 origins were derived from pBb(RK2)1k-GFPuv (8), while the gentamicin resistance cassette was derived from pMQ30 (19), and the spectinomycin cassette was derived from pSR43.6 (20). The RK2 vectors do not contain the *mob* element. A full description of each pGinger vector can be found in **Table 1**.

**Table 1:** Plasmids in the pGinger suite: Relevant characteristics of pGinger plasmids including origin of replication, antibiotic selection, promoter characteristics, and if applicable inducing molecule. JBEI public registry numbers are also included for digital accessibility.

| Name | Origin | Marker | Promoter Class | System | Inducer | JBEI ICE No. |
| --- | --- | --- | --- | --- | --- | --- |
| pGingerBK-J23100 | BBR1 | Kanamycin | Constitutive | J23100 | NA | JPUB_020797 |
| pGingerBK-J23101 | BBR1 | Kanamycin | Constitutive | J23101 | NA | JPUB_020799 |
| pGingerBK-J23102 | BBR1 | Kanamycin | Constitutive | J23102 | NA | JPUB_020815 |
| pGingerBK-J23103 | BBR1 | Kanamycin | Constitutive | J23103 | NA | JPUB_020801 |
| pGingerBK-J23104 | BBR1 | Kanamycin | Constitutive | J23104 | NA | JPUB_020803 |
| pGingerBK-J23105 | BBR1 | Kanamycin | Constitutive | J23105 | NA | JPUB_020817 |
| pGingerBK-J23106 | BBR1 | Kanamycin | Constitutive | J23106 | NA | JPUB_020793 |
| pGingerBK-J23107 | BBR1 | Kanamycin | Constitutive | J23107 | NA | JPUB_020819 |
| pGingerBK-J23108 | BBR1 | Kanamycin | Constitutive | J23108 | NA | JPUB_020821 |
| pGingerBK-J23110 | BBR1 | Kanamycin | Constitutive | J23110 | NA | JPUB_020805 |
| pGingerBK-J23111 | BBR1 | Kanamycin | Constitutive | J23111 | NA | JPUB_020807 |
| pGingerBK-J23113 | BBR1 | Kanamycin | Constitutive | J23113 | NA | JPUB_020809 |
| pGingerBK-J23114 | BBR1 | Kanamycin | Constitutive | J13114 | NA | JPUB_020811 |
| pGingerBK-J23117 | BBR1 | Kanamycin | Constitutive | J13117 | NA | JPUB_020795 |
| pGingerBK-J23118 | BBR1 | Kanamycin | Constitutive | J23118 | NA | JPUB_020823 |
| pGingerBK-J23119 | BBR1 | Kanamycin | Constitutive | J23119 | NA | JPUB_020813 |
| pGingerBK-JE | BBR1 | Kanamycin | Inducible | Jungle Express | Crystal Violet | JPUB_020825 |
| pGingerBK-NahR | BBR1 | Kanamycin | Inducible | Psal/NahR | Salicylic acid | JPUB_020831 |
| pGingerBK-RhaS | BBR1 | Kanamycin | Inducible | Prha/RhaS | Rhamnose | JPUB_020829 |
| pGingerBK-TetR | BBR1 | Kanamycin | Inducible | Ptet/TetR | Oxytetracycline | JPUB_020835 |
| pGingerBK-XylS | BBR1 | Kanamycin | Inducible | Pm/XylS | Benzoate | JPUB_020827 |
| pGingerBK-Lac | BBR1 | Kanamycin | Inducible | LacO1/LacI | IPTG | JPUB_020833 |
| pGingerBK-LacUV5 | BBR1 | Kanamycin | Inducible | LacUV5/LacI | IPTG | JPUB_020837 |
| pGingerBG-JE | BBR1 | Gentamicin | Inducible | Jungle Express | Crystal Violet | JPUB_020847 |
| pGingerBS-JE | BBR1 | Spectinomycin | Inducible | Jungle Express | Crystal Violet | JPUB_020855 |
| pGingerRK-JE | RK2 | Kanamycin | Inducible | Jungle Express | Crystal Violet | JPUB_020871 |
| pGingerRG-JE | RK2 | Gentamicin | Inducible | Jungle Express | Crystal Violet | JPUB_020881 |
| pGingerRS-JE | RK2 | Spectinomycin | Inducible | Jungle Express | Crystal Violet | JPUB_020863 |
| pGingerBG-NahR | BBR1 | Gentamicin | Inducible | Psal/NahR | Salicylic acid | JPUB_020845 |
| pGingerBS-NahR | BBR1 | Spectinomycin | Inducible | Psal/NahR | Salicylic acid | JPUB_020853 |
| pGingerRK-NahR | RK2 | Kanamycin | Inducible | Psal/NahR | Salicylic acid | JPUB_020869 |
| pGingerRG-NahR | RK2 | Gentamicin | Inducible | Psal/NahR | Salicylic acid | JPUB_020879 |
| pGingerRS-NahR | RK2 | Spectinomycin | Inducible | Psal/NahR | Salicylic acid | JPUB_020859 |
| pGingerBG-TetR | BBR1 | Gentamicin | Inducible | Ptet/TetR | Oxytetracycline | JPUB_020843 |
| pGingerBS-TetR | BBR1 | Spectinomycin | Inducible | Ptet/TetR | Oxytetracycline | JPUB_020851 |
| pGingerRK-TetR | RK2 | Kanamycin | Inducible | Ptet/TetR | Oxytetracycline | JPUB_020865 |
| pGingerRG-TetR | RK2 | Gentamicin | Inducible | Ptet/TetR | Oxytetracycline | JPUB_020877 |
| pGingerRS-TetR | RK2 | Spectinomycin | Inducible | Ptet/TetR | Oxytetracycline | JPUB_020861 |
| pGingerBG-LacO1 | BBR1 | Gentamicin | Inducible | LacO1/LacI | IPTG | JPUB_020841 |
| pGingerBS-Lac | BBR1 | Spectinomycin | Inducible | LacO1/LacI | IPTG | JPUB_020849 |
| pGingerRK-LacO1 | RK2 | Kanamycin | Inducible | LacO1/LacI | IPTG | JPUB_020867 |
| pGingerRG-LacO1 | RK2 | Gentamicin | Inducible | LacO1/LacI | IPTG | JPUB_020875 |
| pGingerRS-LacO1 | RK2 | Spectinomycin | Inducible | LacO1/LacI | IPTG | JPUB_020857 |

### Evaluation of Constitutive Expression pGinger Plasmids

To evaluate the relative strength of constitutive Anderson promoters in the context of the pGinger vectors, plasmids were introduced into both *P. putida* and *E. coli*. Fluorescence was measured after growth in LB medium after 24 hours. When fluorescence was normalized to cell density, expression from Anderson promoters showed significant correlation (Spearman’s ρ = 0.49, *p* = 0.045) between *P. putida* and *E. coli* (**Figure 2**). Promoters J23103 and J23113 were significantly stronger in *E. coli* than in *P. putida*, while promoter J23111 was significantly stronger in *P. putida*. Promoter sequences and mean expression values in both *E. coli* and *P. putida* are listed in **Table 2**.

**Figure 2:**
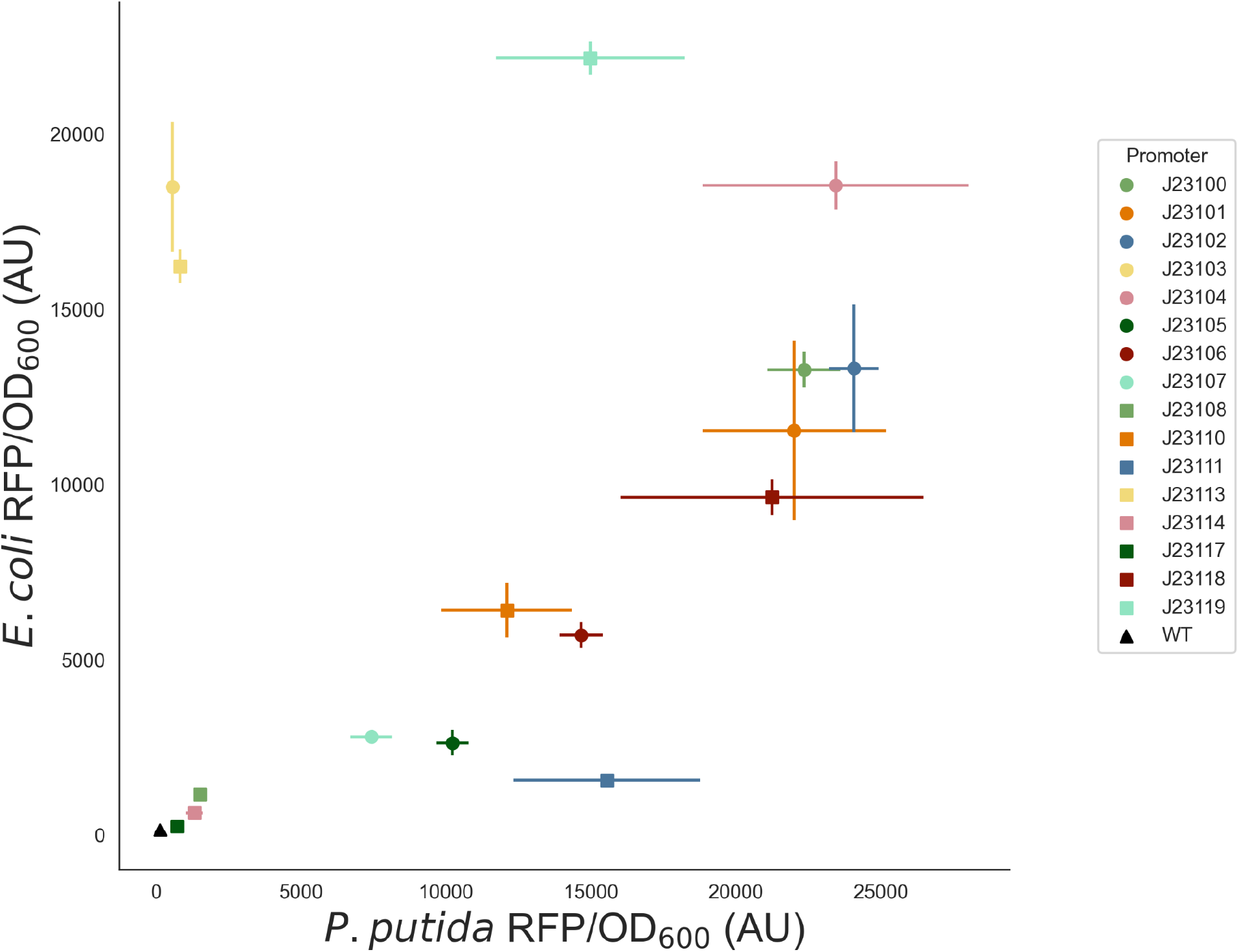
Activity of Constitutive Promoters in *E. coli* and *P. putida*. RFP expression normalized to cell density from Anderson promoters within either *E. coli* (y-axis) or *P. putida* (x-axis) are shown with standard deviations ( n=3). The background fluorescence of the two bacteria is indicated by WT (wild-type). Optical density measurements are shown in Figure S1.

**Table 2:**
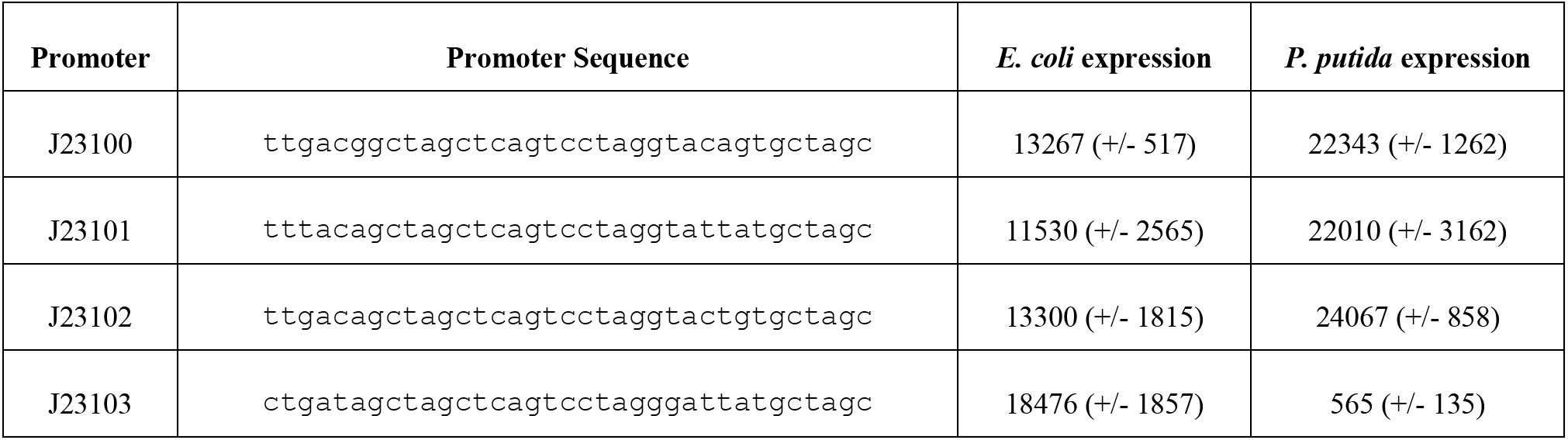
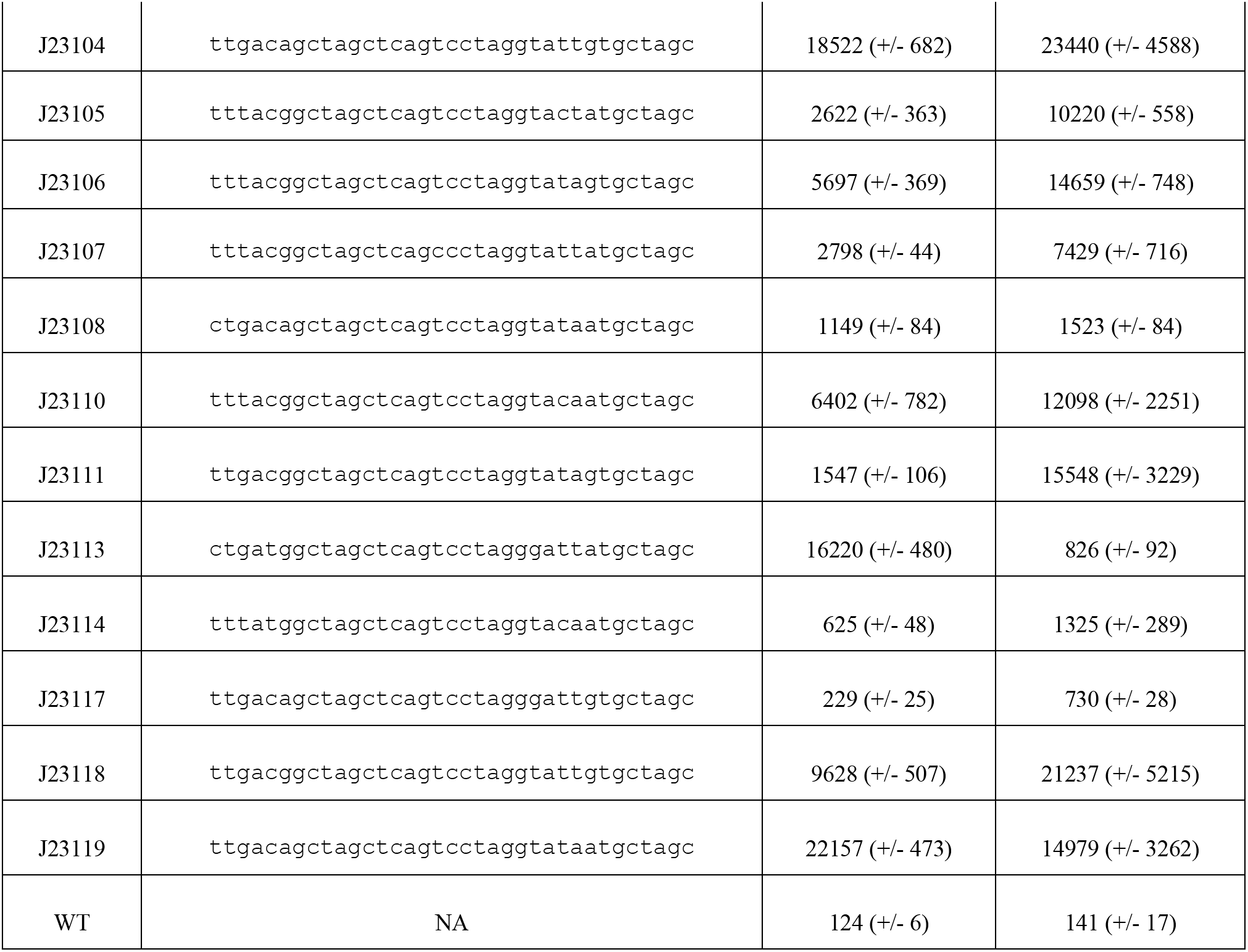
Expression of pGinger Anderson promoters: For each Anderson promoter the sequence is provided as well as the mean cell density normalized RFP fluorescence in both *E. coli* and *P. putida*. Standard deviations are provided in parentheses, n=3. The background fluorescence of E. coli is indicated by WT (wild-type).

### Evaluation of Inducible pGinger Plasmids

The expression of the seven inducible systems within the pGinger suite was evaluated using the BBR1 origin and kanamycin marker (pGingerBK) against a titration of the inducer in both *E. coli* and *P. putida* (**Figure 3**). All systems showed inducibility in *E. coli*, and all but the rhamnose inducible system Prha/RhaS showed inducibility in *P. putida*. Relevant expression characteristics of the inducible pGingerBK vectors in both tested bacteria are listed in **Table 3**. The strongest normalized expression from an inducible system in *E. coli* was the Ptet/TetR system, while both the strongest in *P. putida* were found to be Psal/NahR and Jungle Express inducible systems, which showed nearly identical maximal expression. In both bacteria, the Jungle Express system demonstrated the greatest level of induction relative to background expression.

**Figure 3:**
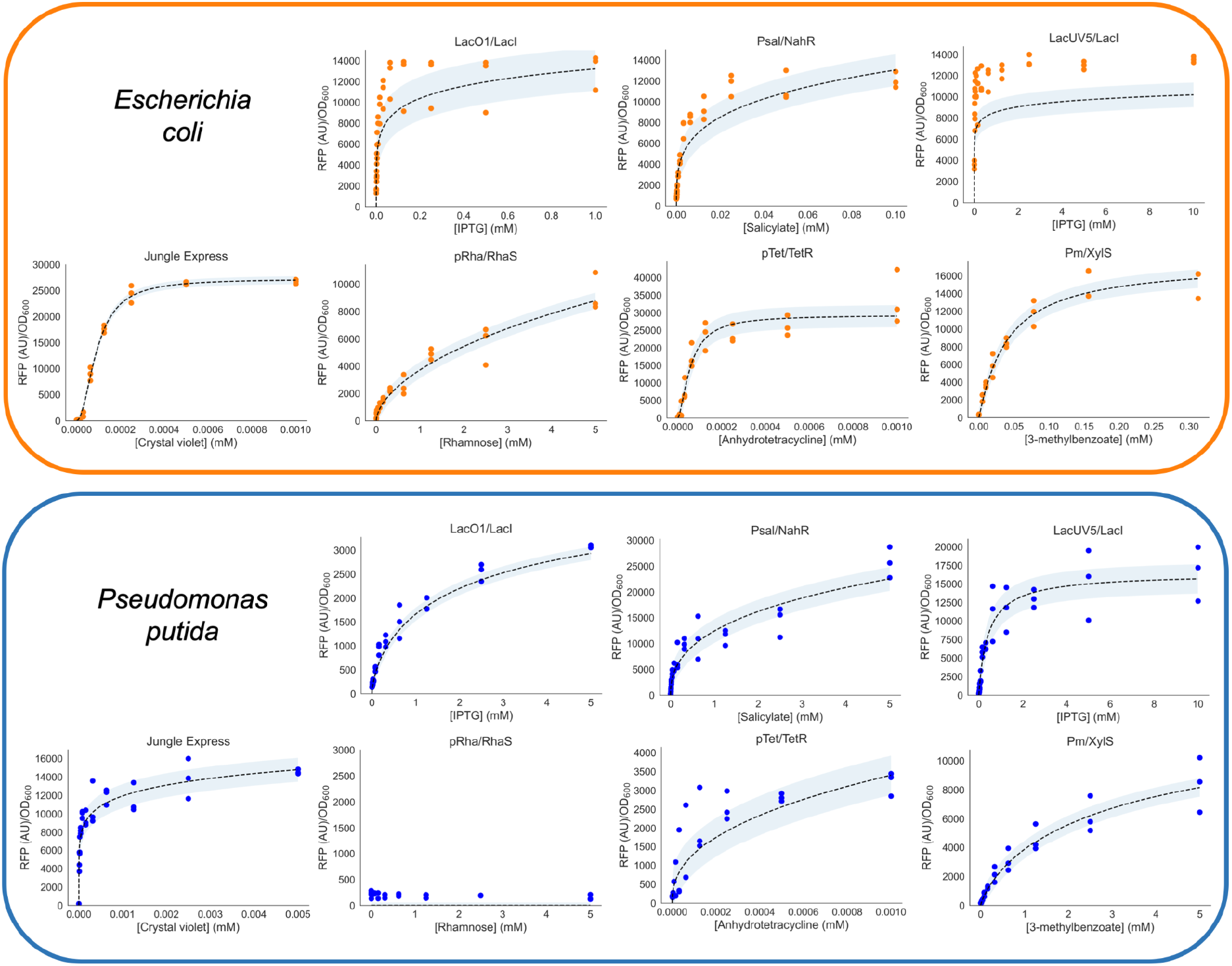
Activity of Inducible Systems in *E. coli* and *P. putida*. RFP expression normalized to cell density (y-axis) from inducible systems within either *E. coli* (top panel in orange) or *P. putida* (bottom panel in blue) as a function of inducer concentration in mM (x-axis). Fits to the Hill equation are shown as dashed lines and shaded to show confidence intervals. Raw data points are overlaid ( n=3). Corresponding optical density measurements are shown in Figure S2.

**Table 3:** Inducible Systems in *E. coli* and *P. putida*: For each inducible system on a BBR1 origin with a kanamycin marker, the experimentally observed background (uninduced) fluorescence and maximal fluorescence are given for both *E. coli* and *P. putida*. Standard deviations are provided in parentheses, n=3. Additionally, the inducer concentration used to achieve maximal expression and the relative induction levels are listed.

| System | Organism | Background | Max Exp. | Max Conc. | Induction |
| --- | --- | --- | --- | --- | --- |
| LacO1/LacI | *E. coli* | 1510 (+/- 186) | 13127 (+/- 1693) | 1 mM | 9x |
|  | *P. putida* | 137 (+/- 5) | 3074 (+/- 30) | 5 mM | 22x |
| Psal/NahR | *E. coli* | 762 (+/- 130) | 12051 (+/- 759) | 100 uM | 16x |
|  | *P. putida* | 237 (+/- 5) | 25697 (+/- 2976) | 5 mM | 108x |
| LacUV5/LacI | *E. coli* | 3577 (+/- 36) | 10137 (+/- 855) | 10 mM | 4x |
|  | *P. putida* | 203 (+/- 53) | 16622 (+/- 3671) | 10 mM | 82x |
| Jungle Express | *E. coli* | 104 (+/- 7) | 26717 (+/- 418) | 1 uM | 257x |
|  | *P. putida* | 176 (+/- 5) | 14552 (+/- 145) | 2.5 uM | 83x |
| Prha/RhaS | *E. coli* | 172 (+/- 42) | 9251 (+/- 1389) | 5 mM | 54x |
|  | *P. putida* | NA | NA | NA | NA |
| Ptet/TetR | *E. coli* | 341 (+/- 10) | 33631 (+/- 7692) | 1 uM | 98x |
|  | *P. putida* | 176 (+/- 9) | 3214 (+/- 319) | 1 uM | 18x |
| Pm/XylS | *E. coli* | 329 (+/- 57) | 15280 (+/- 1590) | 313 uM | 46x |
|  | *P. putida* | 161 (+/- 7) | 8401 (+/- 1877) | 5 mM | 52x |

To evaluate the effect of varying origin and selectable markers on expression from inducible systems, all six variants of the Jungle Express, LacO1/LacI, Psal/NahR, and Ptet/TetR were investigated for their dose-response to their inducer molecules in *E. coli* (**Figure 4**). Relevant expression parameters are listed in **Table 4**. In general, BBR1 variants showed greater expression than RK2 origin plasmids, which is expected given the higher copy of BBR1 plasmids in *E. coli* (11). Amongst the pGinger Jungle Express vectors, both pGingerRS-JE and pGingerRK-JE showed dose-responses distinct from the other vectors (**Figure 4A**). Notably, all pGingerRS (RK2-Spectinomycin) plasmids showed the lowest expression across each system tested (**Table 4**).

**Figure 4:**
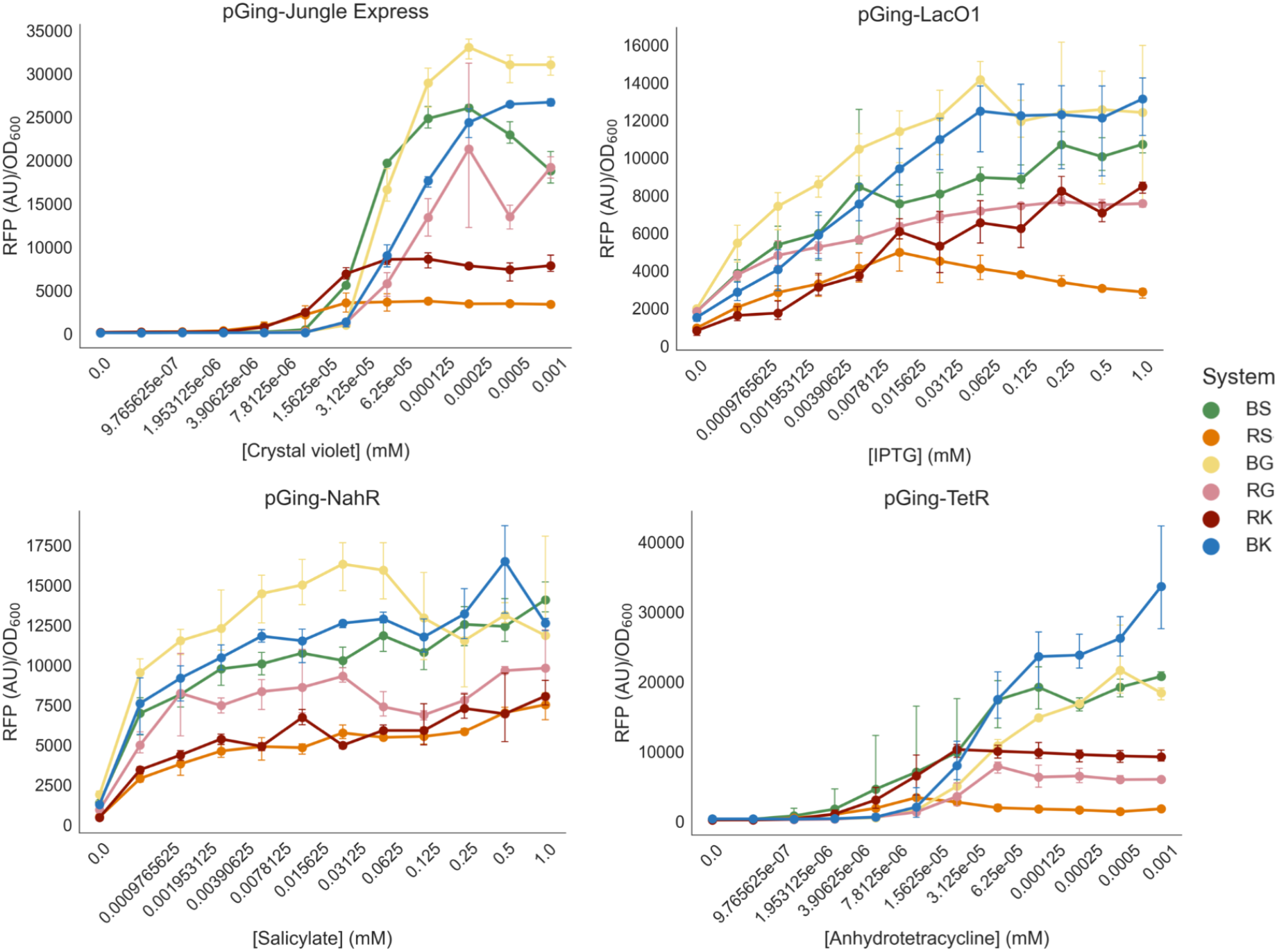
Activity of Inducible pGinger variants in *E. coli*. For origin and selection marker pGinger variants of Jungle Express (A), LacO1/LacI (B), Psal/NahR (C), and pTet/TetR (D), dose-response curves of normalized RFP expression are shown as a function of mM inducer. Error bars represent standard deviations (n=3). Note that the x-axis is non-linear. Corresponding optical density measurements are shown in Figure S3. Kinetic experiments for these systems are shown in Figure S4-S7.

**Table 4:** Inducible pGinger variants in *E. coli*: For pGinger variants of LacO1/LacI, Psal/NahR, Jungle Express, and Ptet/TetR inducible systems, the experimentally observed background (uninduced) fluorescence and maximal fluorescence in *E. coli* are provided. Standard deviations are provided in parentheses, n=3. Additionally, the inducer concentration used to achieve maximal expression and the relative induction levels are listed.

| System | Origin | Marker | Background | Max Exp. | Max Conc. | Induction |
| --- | --- | --- | --- | --- | --- | --- |
| LacO1/LacI | BBR1 | Kan | 1510 (+/- 186) | 13127 (+/- 1693) | 1 mM | 9x |
| LacO1/LacI | BBR1 | Gent | 1976 (+/- 70) | 14143 (+/- 1608) | 63 uM | 7x |
| LacO1/LacI | BBR1 | Spec | 1830 (+/- 320) | 10694 (+/- 943) | 250 uM | 6x |
| LacO1/LacI | RK2 | Kan | 803 (+/- 216) | 8213 (+/- 721) | 250 uM | 10x |
| LacO1/LacI | RK2 | Gent | 1823 (+/- 78) | 7654 (+/- 230) | 250 uM | 4x |
| LacO1/LacI | RK2 | Spec | 950 (+/- 46) | 4967 (+/- 895) | 16 uM | 5x |
| Psal/NahR | BBR1 | Kan | 1260 (+/- 57) | 16484 (+/- 1693) | 500 uM | 9x |
| Psal/NahR | BBR1 | Gent | 1877 (+/- 334) | 16317 (+/- 1534) | 31 uM | 9x |
| Psal/NahR | BBR1 | Spec | 1372 (+/- 84) | 14083 (+/- 984) | 1 mM | 10x |
| Psal/NahR | RK2 | Kan | 445 (+/- 39) | 8048 (+/- 859) | 1 mM | 18x |
| Psal/NahR | RK2 | Gent | 934 (+/- 56) | 9299 (+/- 460) | 1 mM | 10x |
| Psal/NahR | RK2 | Spec | 482 (+/- 69) | 7510 (+/- 817) | 1 mM | 16x |
| Jungle Express | BBR1 | Kan | 104 (+/- 6) | 267175 (+/- 418) | 1 uM | 257x |
| Jungle Express | BBR1 | Gent | 136 (+/- 19) | 33060(+/- 1185) | 250 nM | 243x |
| Jungle Express | BBR1 | Spec | 152 (+/- 5) | 26039 (+/- 355) | 250 nM | 171x |
| Jungle Express | RK2 | Kan | 163 (+/- 27) | 8616 (+/- 902) | 125 nM | 52x |
| Jungle Express | RK2 | Gent | 138 (+/- 51) | 21314 (+/- 9517) | 250 nM | 155x |
| Jungle Express | RK2 | Spec | 168 (+/- 5) | 3763 (+/- 204) | 125 nM | 22x |
| Ptet/TetR | BBR1 | Kan | 341 (+/- 10) | 33631 (+/- 7692) | 1 uM | 98x |
| Ptet/TetR | BBR1 | Gent | 351 (+/- 32) | 21646 (+/- 5579) | 500 nM | 62x |
| Ptet/TetR | BBR1 | Spec | 232 (+/- 43) | 19224 (+/- 3027) | 125 nM | 83x |
| Ptet/TetR | RK2 | Kan | 184 (+/- 5) | 10281 (+/- 967) | 31 nM | 56x |
| Ptet/TetR | RK2 | Gent | 232 (+/- 39) | 7883 (+/- 865) | 63 nM | 34x |
| Ptet/TetR | RK2 | Spec | 197 (+/- 5) | 3399 (+/- 143) | 16 nM | 17x |

## Discussion

The pGinger suite of plasmids offers researchers an array of small, pre-assembled vectors that will permit rapid identification of useful genetic elements in diverse gram-negative bacteria due to the use of broad host-range origins (RK2) and selectable markers known to work across many species (kanamycin, spectinomycin, gentamicin). The compatibility of RK2 and BBR1 origins may also permit researchers to introduce multiple pGinger vectors into a single strain simultaneously (14). In combination with other recent plasmid suites that have been publicly released, the pGinger plasmids have the potential to facilitate more advanced synthetic biology and metabolic engineering efforts in bacterial species that have been traditionally understudied.

## Materials & Methods

### Strains and Media

Cultures were grown in lysogeny broth (LB) Miller medium (BD Biosciences, USA) at 37 °C for *E. coli* XL1-Blue (QB3 Macrolab, USA) and 30 °C for *P. putida* KT2440 (ATCC 47054). The medium was supplemented with kanamycin (50 mg/L, Sigma Aldrich, USA), gentamicin (30 mg/L, Fisher Scientific, USA), or spectinomycin (100mg/L, Sigma Aldrich, USA), when indicated. All other compounds were purchased through Sigma Aldrich (Sigma Aldrich, USA).

### Plasmid Design and Construction

All plasmids were designed using Device Editor and Vector Editor software, while all primers used for the construction of plasmids were designed using j5 software (15, 21, 22). Plasmids were assembled via Gibson Assembly using standard protocols (23). Plasmids were routinely isolated using the Qiaprep Spin Miniprep kit (Qiagen, USA), and all primers were purchased from Integrated DNA Technologies (IDT, Coralville, IA). The fluorescent protein used in all plasmids was mRFP1 (24).

### Plasmid and Sequence Availability

All strains and plasmid sequences from Table 1 can be found via the following link to the JBEI Public Registry: https://public-registry.jbei.org/folders/771. Users can request strains via a MTA.

### Characterization assays

To characterize RFP expression from these vectors, we measured optical density and fluorescence after growth in 96 well plates for 24 hours. First, overnight cultures were inoculated into 5 mL of LB medium from single colonies and grown at 30 °C or 37 °C. These cultures were then diluted 1:100 into 500 μL of LB medium with the appropriate antibiotic in 96 square v-bottom deep well plates (Biotix™ DP22009CVS). For characterization of the inducible systems, inducer was added to wells in the first column of the plate at the maximum concentration tested and diluted two-fold across the plate until the last column, which was left as the zero-inducer control. Plates were sealed with a gas-permeable microplate adhesive film (Axygen™ BF400S) and grown for 24 hours at either 30 °C or 37 °C with shaking at 200 rpm. Optical density was measured at 600 nm, and fluorescence was measured at an excitation wavelength of 535 nm and an emission wavelength of 620 nm. All data was analyzed and visualized using custom Python scripts using the SciPy (25), NumPy (26), Pandas, Matplotlib, and Seaborn libraries. Fits to the Hill equation were done as previously described (27).

## Acknowledgements

The order of co-first authors was determined by who has the reddest hair. Mitchell Thompson is a Simons Foundation Awardee of the Life Sciences Research Foundation. Lucas Waldburger is funded by the National Science Foundation Graduate Research Fellowship. This work was part of the DOE Joint BioEnergy Institute (https://www.jbei.org) supported by the U.S. Department of Energy, Office of Science, Office of Biological and Environmental Research, supported by the U.S. Department of Energy, Energy Efficiency and Renewable Energy, Bioenergy Technologies Office, through contract DE-AC02-05CH11231 between Lawrence Berkeley National Laboratory and the U.S. Department of Energy. The views and opinions of the authors expressed herein do not necessarily state or reflect those of the United States Government or any agency thereof. Neither the United States Government nor any agency thereof, nor any of their employees, makes any warranty, expressed or implied, or assumes any legal liability or responsibility for the accuracy, completeness, or usefulness of any information, apparatus, product, or process disclosed, or represents that its use would not infringe privately owned rights. The United States Government retains and the publisher, by accepting the article for publication, acknowledges that the United States Government retains a nonexclusive, paid-up, irrevocable, worldwide license to publish or reproduce the published form of this manuscript, or allow others to do so, for United States Government purposes. The Department of Energy will provide public access to these results of federally sponsored research in accordance with the DOE Public Access Plan (http://energy.gov/downloads/doe-public-access-plan).

## Contributions

Conceptualization; A.N.P., M.G.T.; Methodology; A.N.P., M.G.T..; Investigation, A.N.P., M.G.T., L.D.K., C.H., K.M.V., L.M.W.; Writing – Original Draft, A.N.P., M.G.T..; Writing – Review and Editing, All authors.; Resources and supervision; P.M.S, J.D.K.

