## Supplemental Material for "The pGinger family of expression plasmids"

**The pGinger family of expression plasmids: Supplemental Material**

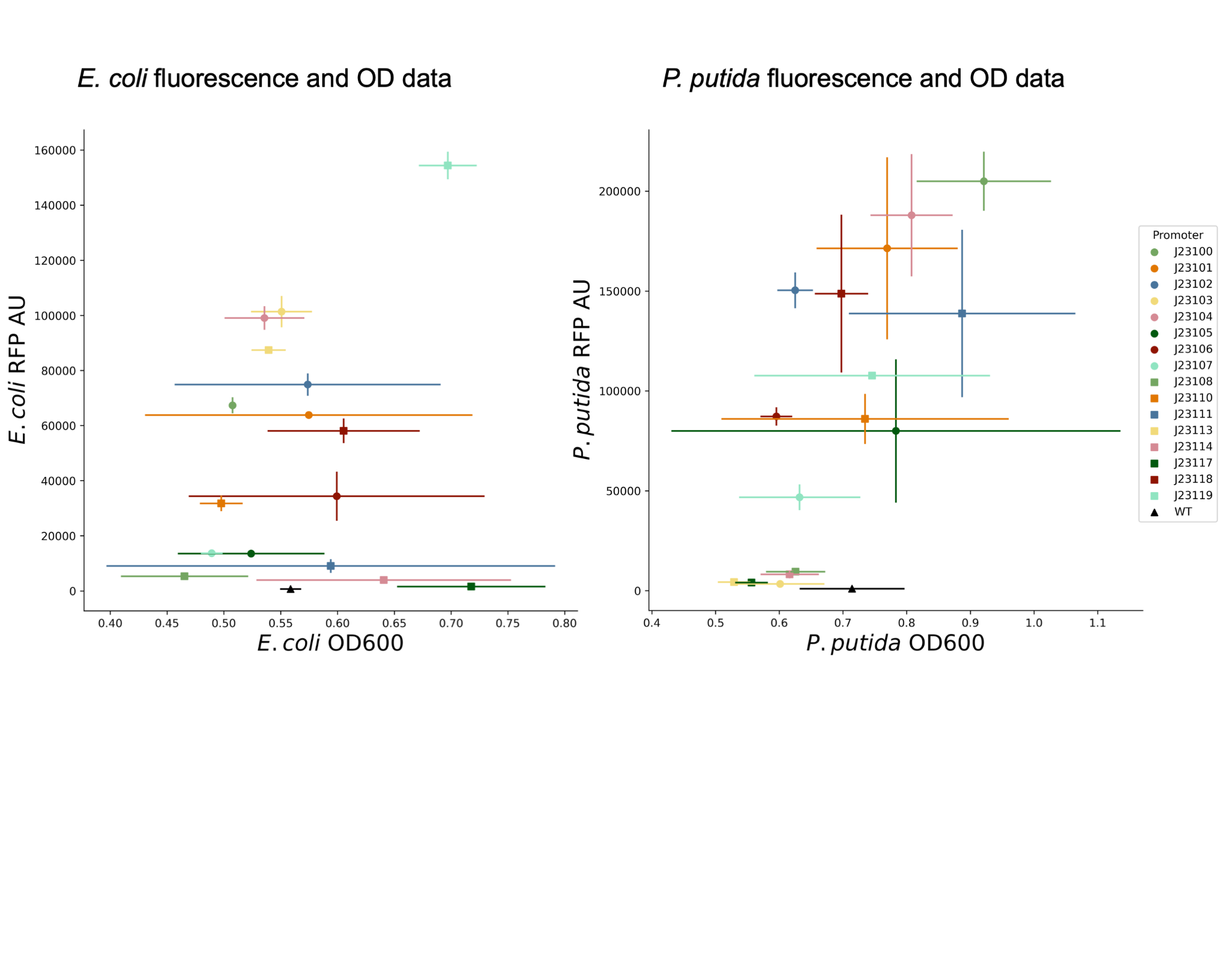
**Figure S1:** Optical density and fluorescence measurements after 24 hours of growth, corresponding to the normalized fluorescence data presented in Figure 2. Error bars represent standard deviation (n=3, error bars = std. dev.)

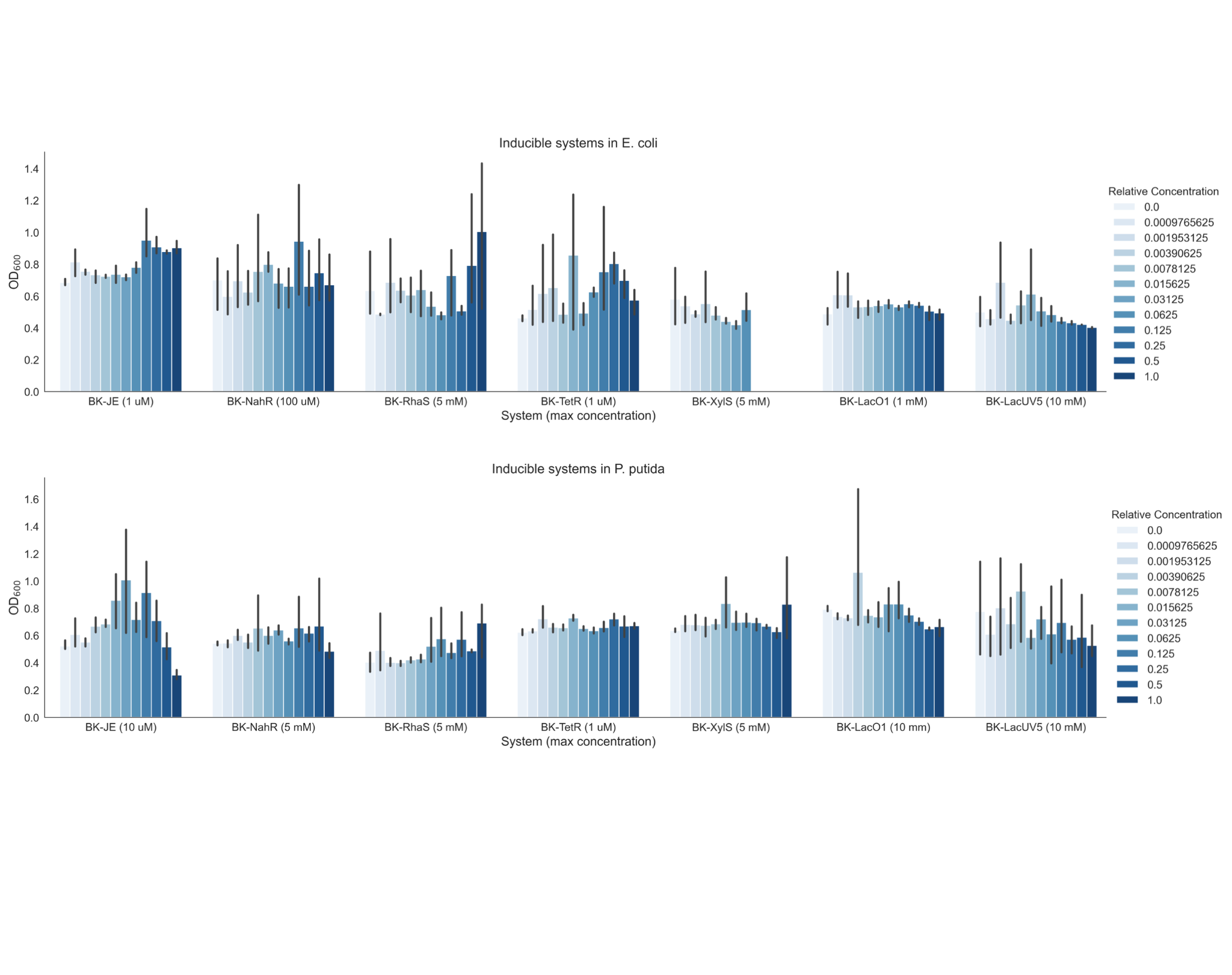
**Figure S2:** Corresponding optical density endpoint measurements for data presented in Figure 3 (n=3, error bars = std. dev.)

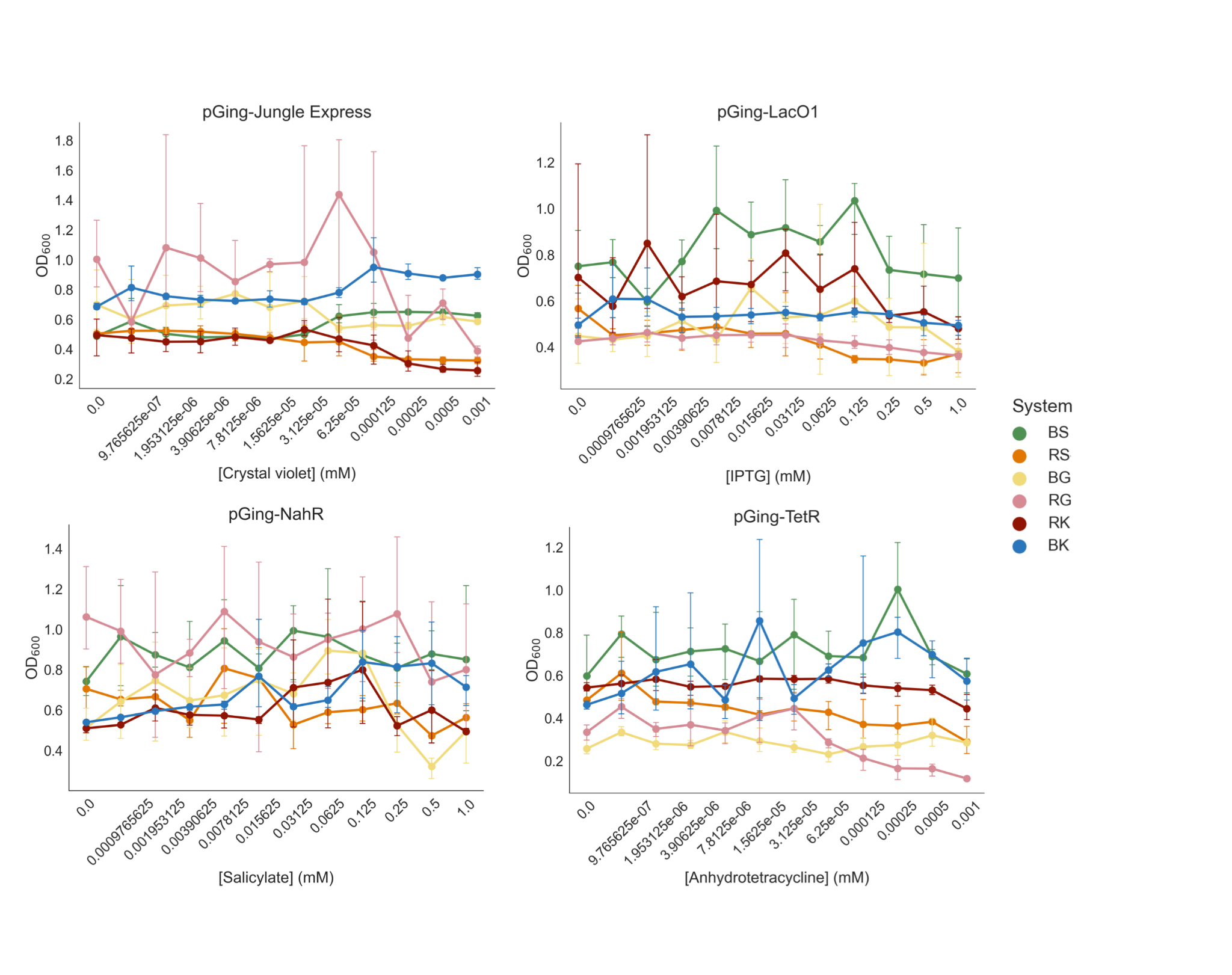
**Figure S3:** Corresponding optical density measurements for data presented in Figure 4 (n=3, error bars = std. dev.)

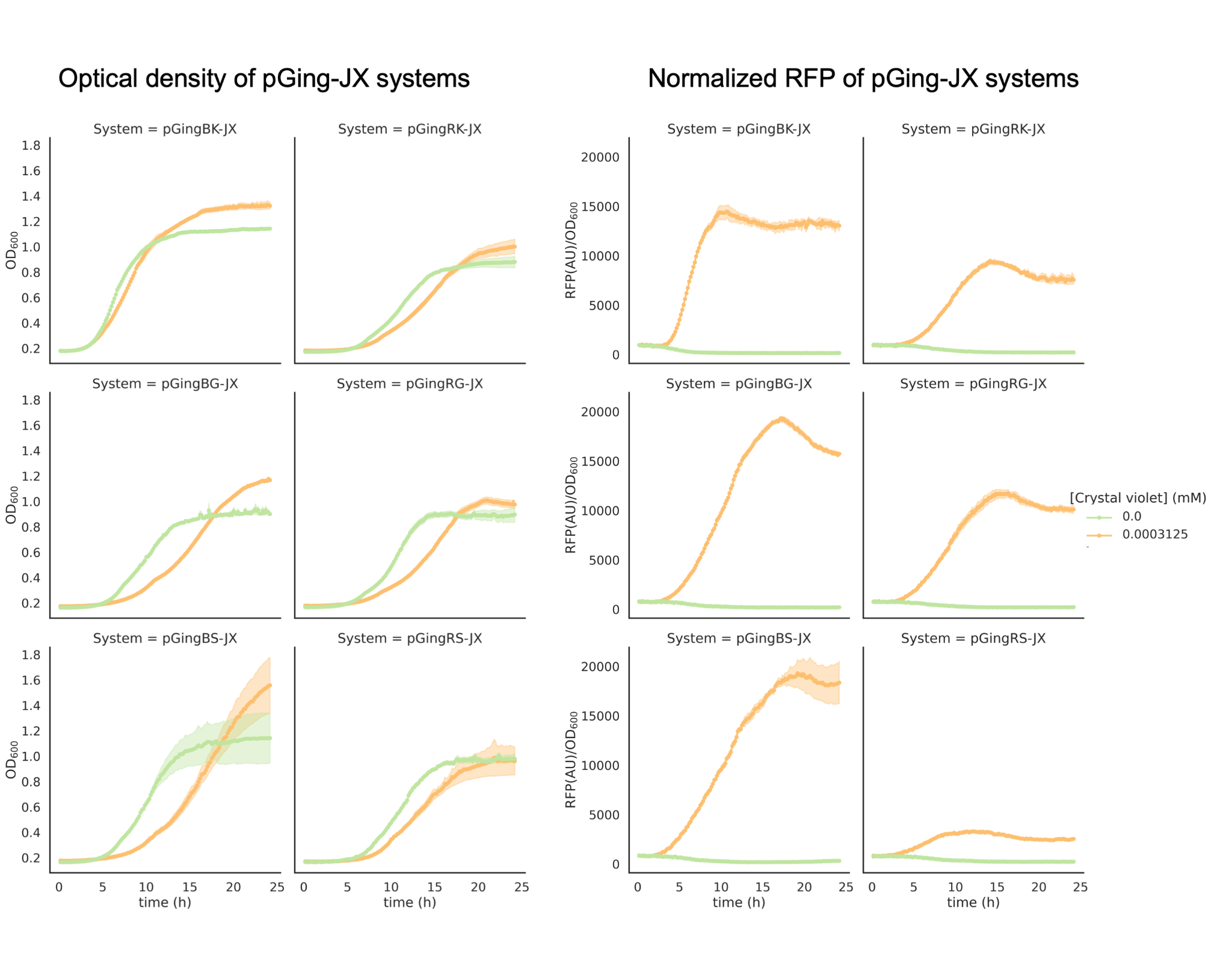
**Figure S4:** Kinetic growth and fluorescence data for pGing-JX systems in *E. coli* (n=3).

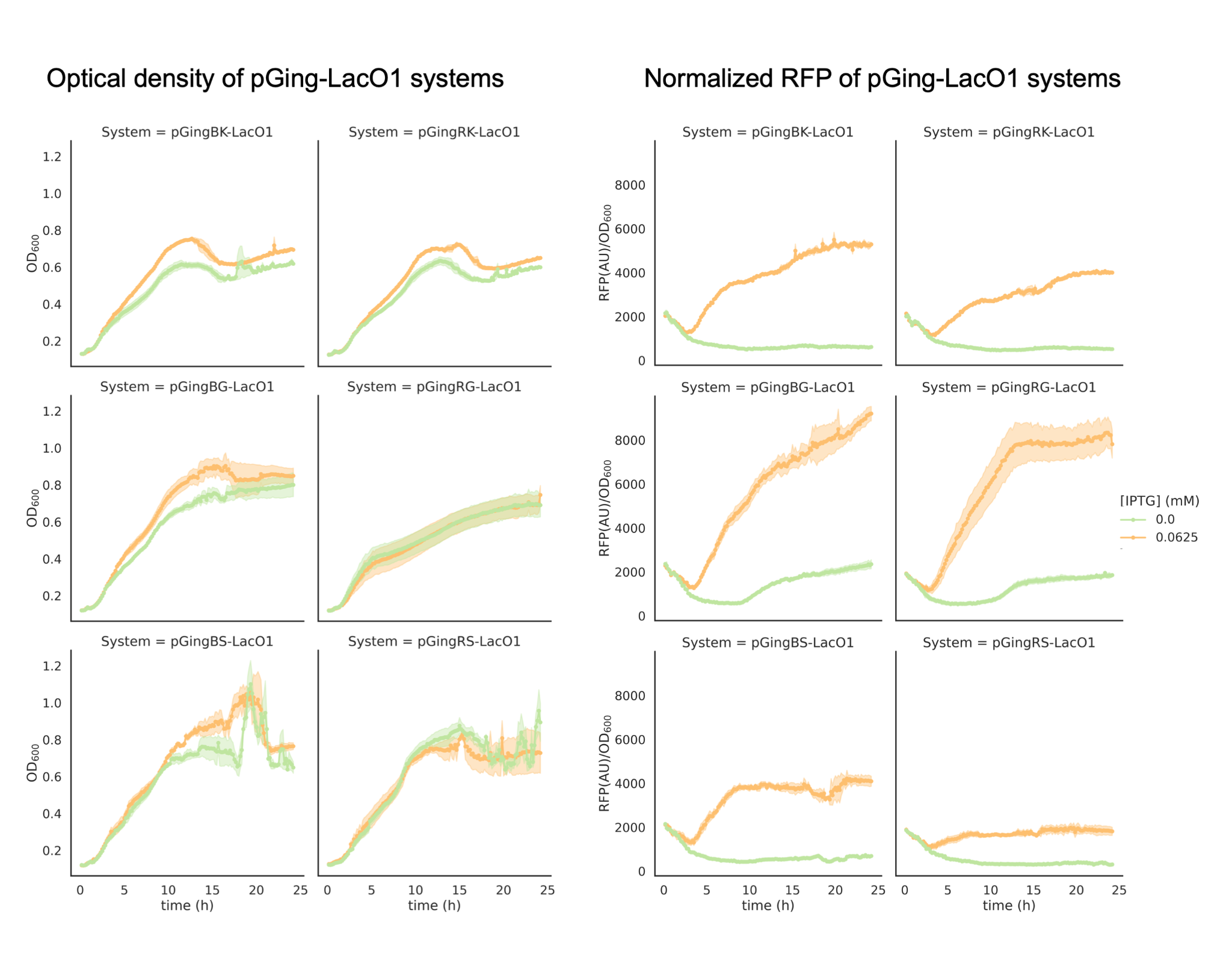
**Figure S5:** Kinetic growth and fluorescence data for pGing-LacO1 systems in *E. coli* (n=3).

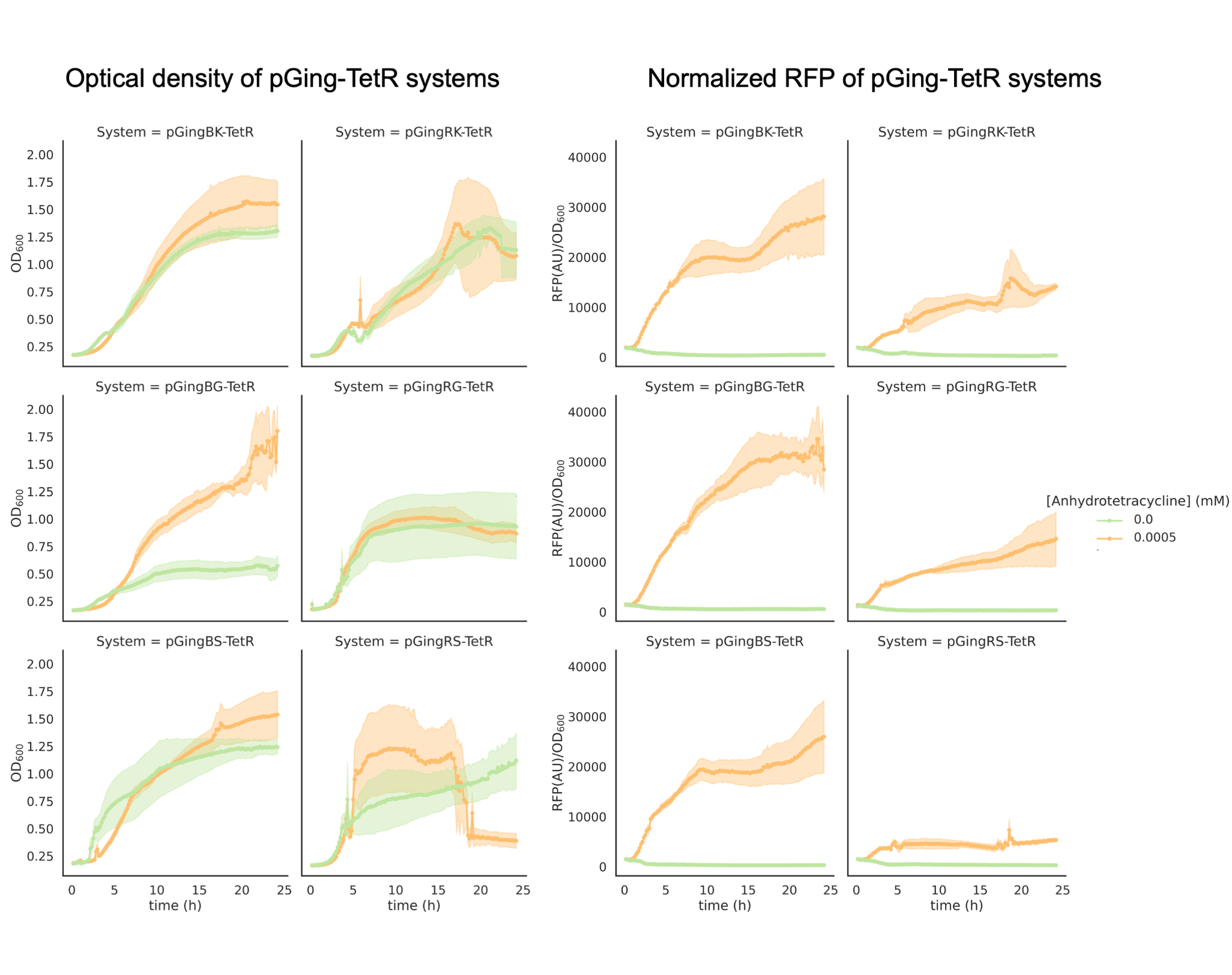
**Figure S6:** Kinetic growth and fluorescence data for pGing-TetR systems in *E. coli* (n=3).

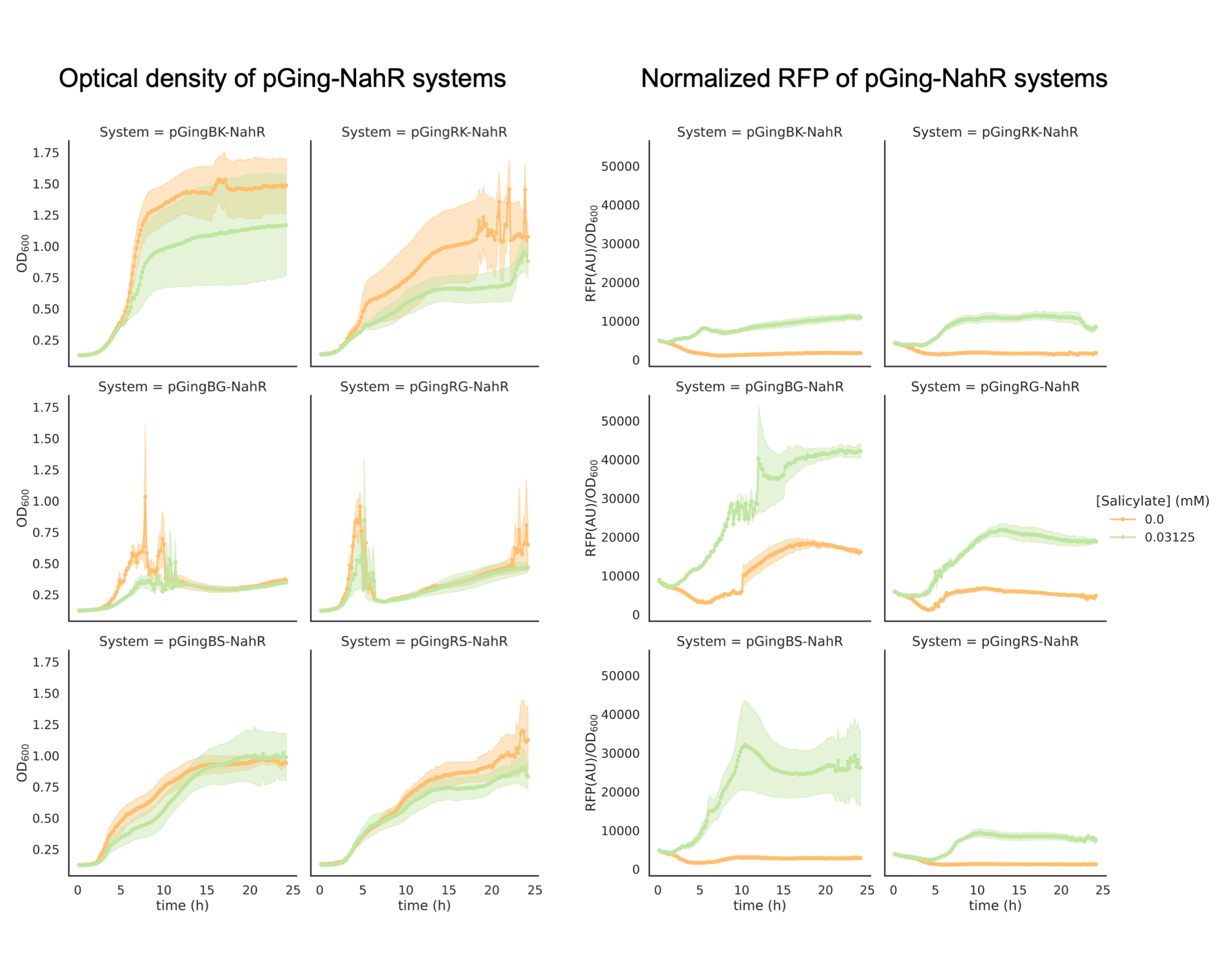
**Figure S7:** Kinetic growth and fluorescence data for pGing-NahR systems in *E. coli* (n=3).
